# Neopep: Reinforcement Learning from Machine-Learned Force Field Energy for Non-Canonical Peptide Structure Prediction

**DOI:** 10.64898/2026.10.22.760799

**Authors:** Dhanya Jagannathan, Philip M. Kim

## Abstract

Peptide therapeutics occupy a compelling middle ground between small molecules and larger proteins, and in practice, most approved peptide drugs incorporate non-canonical amino acids (ncAAs) or other non-peptidic chemistries to elicit improved physiochemical properties (Yaseen Khan et al. (2026)). The chemical space of known ncAAs numbers in the thousands, yet current state-of-the-art sequence-to-structure prediction models handle them poorly, because fewer than 0.02% of residues deposited in the Protein Data Bank (PDB) (Berman et al. (2003)), the main source of training data, correspond to ncAAs. Our central idea is that this data bottleneck can be overcome by coupling generative structure prediction with reinforcement learning (RL) from machine-learned force field (MLFF)-derived energy. We present **Neopep**, a sequence-conditioned diffusion model for ncAA-containing peptide structure prediction, first pretrained on PDB structures to establish a structural prior and then fine-tuned via RL: for ncAA-containing peptide sequences, the model proposes structures that are scored by a frozen MLFF, and this energy signal is backpropagated directly into the model to progressively lower the energy of its predictions. MLFFs have demonstrated near-quantum accuracy across much of chemical space, approacing DFT-level accuracy at a fraction of the computational cost (Kabylda et al. (2025)). This makes them substantially more accurate than classical, empirically derived force fields, which are parametrised for the canonical 20 amino acids and do not generalise to arbitrary chemistries. While Neopep is trained on a fixed set of 189 ncAAs, the underlying architecture operates at atom-level granularity and is in principle generalisable to any chemistry. We show that our model is able to produce structures of ncAA-containing peptides that achieve lower RMSD values on average to the ground truth structures, in comparison to without finetuning and other state-of-the-art structure prediction models.

## 1 Introduction

Peptide therapeutics are a compelling middle ground between small molecules and larger proteins. Relative to small molecules, they offer greater target specificity and lower toxicity, and relative to larger proteins, they are more stable and less costly to produce (Abdin & Kim (2024))(Wang et al. (2022)). However, the canonical vocabulary of 20 proteinogenic amino acids constrains the chemical space available for designing such peptide therapeutics. In practice, most approved peptide drugs incorporate ncAAs or non-peptidic chemistries for improved physiochemical properties (Yaseen Khan et al. (2026)). Expanding the canonical 20 amino acid vocabulary to the thousands of known non-canonical amino acids (ncAAs)-many of which occur naturally as metabolic intermediates or post-translational modifications, while others are purely synthetic-would allow for the design of novel peptide drugs with unique and improved properties (Lee & Kim (2026))(Budisa (2025)). Realizing this potential from a computational perspective incentivizes the need to accurately model the structures of peptides containing ncAAs. Current state-of-the-art (SOTA) sequence-to-structure prediction models (Abramson et al. (2024))(Discovery et al. (2024))(Passaro et al. (2025)) are able to accept ncAAs in the input sequence, but are unable to model their structures well. This is due to the fact that the Protein Data Bank (PDB), the main source of training data for these models, has very little peptides with ncAAs as less than 0.02% of the residues deposited in the PDB correspond to ncAAs (Lee & Kim (2026)). The central question then becomes, how can we navigate this data scarcity concern to effectively model peptides with ncAAs?

To address this, we turn to the physical energy function as additional training signal. This is grounded in the Boltzmann distribution, which states that: for a particular sequence, low-energy structures are more probable than high-energy ones, meaning that a peptide’s native structure is a low energy one. Concretely, instead of training a generative sequence-to-structure prediction model only to reproduce the ground-truth structures we happen to have, we can train it to produce structures that are low in energy-which is our way out of the data problem. Where structural data is absent, physics is not. We obtain this oracle energy from a machine-learned force field (Unke et al. (2021)), which are trained on large quantum-chemical datasets, and now reach near-quantum accuracy across the chemical space of organic and biological molecules, with speed much faster than traditional Density Functional Theory (DFT) (Kabylda et al. (2025)). Additionally, MLFFs are not restricted to the 20 proteinogenic amino acids like classical force fields are - as they function at the atomic level and do not require residue-level parametrization. Specifically, we use SO3LR (Kabylda et al. (2025)), built on the SO3krates architecture (Frank et al. (2024)) and trained on four million molecular complexes at the PBE0+MBD level of theory. Other potentials such as MACE-OFF (Kovács et al. (2025)), AIMNet (Zubatyuk et al. (2019)) (Anstine et al. (2024)), AI2BMD (Wang et al. (2024)) are also available, however, we choose SO3LR for our purposes because of its explicit treatment of molecular charges, and thorough analyses that demonstrate its effectiveness on bio-molecular systems specifically.

### Concretely, our contribution is as follows

We introduce **Neopep**, a sequence-conditioned diffusion model to predict structures of peptides with ncAAs. We first pretrain a sequence-conditioned structure predictor following the AlphaFold3 (AF3) architecture on structures from the PDB, which we subsequently finetune with an energy reward from a MLFF, following the Direct Reward Fine-Tuning (DRaFT) framework (Clark et al. (2023)).

## 2 Related Work

### Physical energy function as training signal

The field of sequence-conditioned protein structure prediction has increasingly shifted from modeling a single static structure to sampling the full conformational ensemble a protein can adopt, aka, sampling from the equilibrium, or Boltzmann distribution. While early progress on this front came from (1) architectural advances, like the shift from AlphaFold2 (AF2) (Jumper et al. (2021)) to the diffusion-based AF3 (Abramson et al. (2024)), alongside models such as Boltz (Passaro et al. (2025)), BioEmu (Lewis et al. (2025)), the Distributional Graphformer (Zheng et al. (2024)), and AlphaFlow (Jing et al. (2024)), and (2) from data-side strategies like incorporation of molecular dynamics trajectories for additional training data (Lewis et al. (2025)), these approaches remain bottlenecked by the need for large volumes of ground-truth structural data. More recently, the field has responded to this with an energy-as-supervision paradigm, in which the physical energy function itself serves as the training signal for equilibrium sampling, in place of ground-truth structures. We see this in the Boltzmann generator framework (Noé et al. (2019)), and a recent application of it to model the equilibrium distribution of peptides with canonical amino acids (Abdin & Kim (2024)), as well as the pre-training scheme introduced in the Distributional Graphformer paper (Zheng et al. (2024)). Additionally, this energy-as-supervision paradigm appears again in the application of GFLowNets to equilibrium small-molecule conformer generation (Volokhova et al. (2024)). To our knowledge, however, these efforts have remained confined to canonical amino acid space, and has not yet been applied to address the data scarcity concerning non-canonical chemistries.

### Modeling and design of proteins with ncAAs

Rarefold (Li et al. (2025)), a structure prediction model for proteins containing ncAAs, extends the AF2 architecture by treating each amino acid, canonical or noncanonical, as a distinct token. This is different from the AF3 archictecture (Abramson et al. (2024)) which represents ncAAs at atom-level granularity and canonicals at residue-level granularity. However, because each ncAA is represented by a dedicated token, RareFold is fundamentally restricted to ncAAs seen during training, that is a set of 29 ncAAs, and cannot generalize to novel ncAA chemistries. Concurrent work, NCFlow (Lee & Kim (2026)), solves a slightly narrower problem: given a fixed protein backbone and a residue position where an ncAA sits, it predicts the 3D conformation of that single ncAA’s atoms, using only the ncAA’s atom types and bond graph as input, making it generalizable to arbitrary ncAAs unseen during training. To compensate for the scarcity of native ncAA data in the PDB, NCFlow is pretrained on small-molecule conformers and protein-ligand complexes before finetuning on ncAA-protein environments, and demonstrates improved structure prediction accuracy over AF3-based methods. Neopep differs from both these approaches in that, we use a physical energy function to guide us towards plausible structures instead of relying on architectural modifications, while also retaining the goal of full sequence-to-structure prediction rather than single-residue. Additionaly, since our architecture, following the AF3 archictecture, operates at atom-level granularity for ncAA representations, it is generalizable in principle.

## 3 Method

Neopep is trained in two phases, illustrated in Figure 1: (1) pretraining on PDB structures to establish a structural prior (Figure 1a), and (2) fine-tuning the pretrained model with a differentiable MLFF-derived energy reward to steer predictions towards physically plausible structures of ncAA-containing peptides (Figure 1b). We describe the architecture first, then each training phase in turn. Throughout, when we refer to AF3 we mean its open-source reimplementation, Boltz (Passaro et al., 2025).

**Figure 1:**
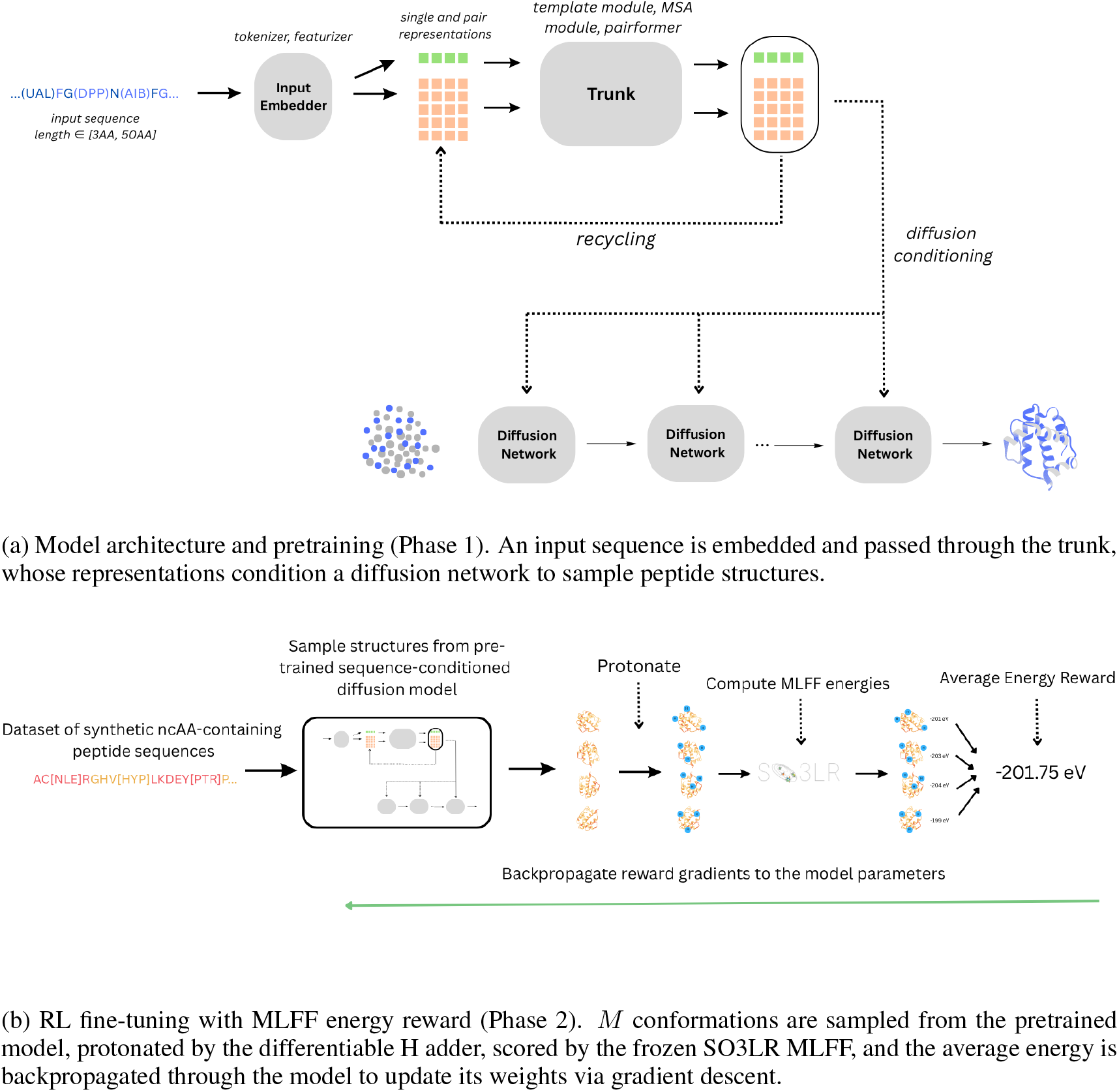
Overview of Neopep. Phase 1 (top) pretrains a Boltz-2-style structure diffusion model on PDB fragments. Phase 2 (bottom) fine-tunes the pretrained model with a differentiable MLFF energy reward via DRaFT-K.

### 3.1 Architecture

As shown in Figure 1a, the Neopep architecture closely follows AF3 (Abramson et al., 2024); we describe it briefly and refer the reader to that paper for full architectural details. The model takes as input a peptide sequence of length 3–50 amino acids, which may contain ncAAs and a cyclic or linear topology flag. The **input embedder** encodes each residue using per-atom features derived from the Chemical Component Dictionary (CCD) (Westbrook et al., 2014). A *pair representation* is constructed alongside a *single representation*, encoding pairwise atom (or residue) relationships. Both representations are refined by the **trunk**, which comprises an MSA module, a template module, a pairformer, and recycling steps (Figure 1a). The refined representations then condition a **diffusion network** that samples plausible all-heavy-atom structures via *T* = 200 denoising steps. Architectural modifications relative to AF3 (reduced MSA and pairformer sizes, disabled template and confidence modules) are detailed in Appendix A.

### 3.2 Training Phase I: Pretraining a Structural Prior

#### Training objective

The pretraining objective follows AF3 and combines a diffusion loss with an auxiliary distogram loss:

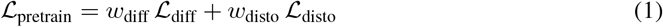

#### Diffusion loss

At each training step the ground-truth heavy-atom coordinates **x**_0_ are corrupted by Gaussian noise at a randomly sampled noise level *σ* following the variance-exploding (VE) schedule: 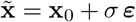, where ***ε*** *∼ N* (**0, I**). The network is trained via **x**-prediction: it predicts the denoised coordinates 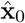 directly, and a weighted MSE loss is computed against **x**_0_.

#### Distogram loss

An auxiliary distogram head is applied to the pair representation from the trunk to predict a discretised distribution over inter-residue distances, supervised with cross-entropy against the ground-truth distances. This loss provides direct supervision on the pair representation independently of the diffusion schedule.

#### Data

Following AF3, we pretrain on PDB structures with a training date cutoff of June 1, 2023; structures deposited after this date form the validation and test sets, which we further filter for sequence similarity with the training set, following AF3 protocol. Because our target application is peptides (3–50 residues), we sample *protein fragments* of this length from longer PDB chains, allowing the full PDB to be used as training data. MSA sequences are capped at 512 and template search is disabled entirely; all other data processing, clustering, and sampling follow Passaro et al. (2025). Additional details are given in Appendix B.

### 3.3 Training Phase II: Fine-Tuning with MLFF-Derived Energy

We then fine-tune the pretrained model from phase 1 to minimize the physical energy of the generated peptide conformations. The objective is as follows:

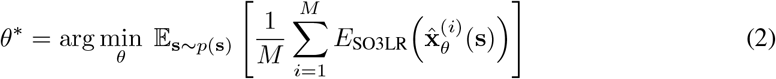

where 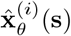 are *M* = 4 conformations sampled from the prior model for peptide sequence **s**, and *E*_SO3LR_ is a frozen MLFF that scores these conformations. Concretely, the sequence **s** is first sampled uniformly, and is then passed as input to the model, to produce proposed structures. These proposals are then scored by the MLFF with an energy value, which is averaged over all M conformations.

#### Algorithm 1

DRaFT-K Fine-Tuning with SO3LR MLFF Energy Reward

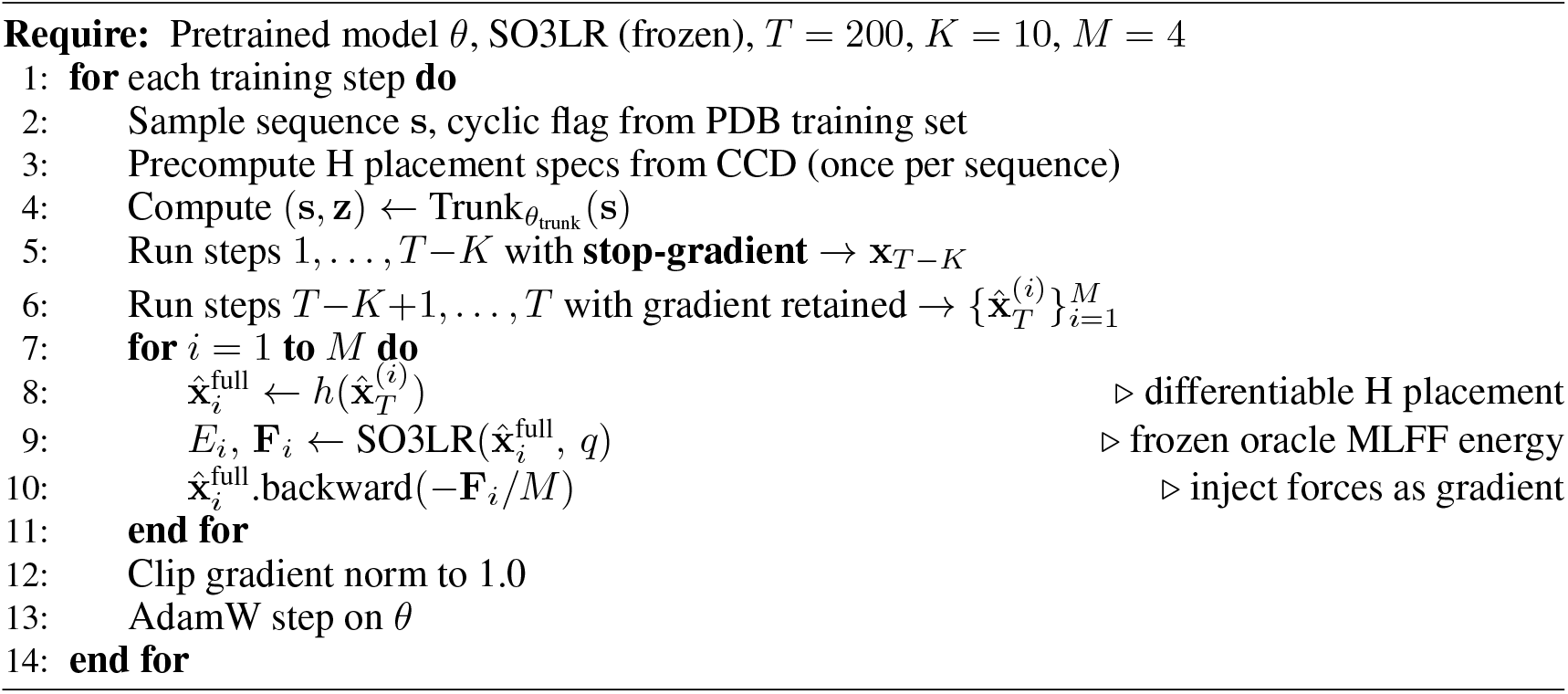

The model is optimized to yield the weights *θ* of the diffusion and trunk networks that best minimize this value. Ablations over the number of conformations *M ∈ {*4, 8, 12, 16} and two energy pooling strategies (average vs. maximum) are reported in Appendix D. In the subsequent sections, we discuss the fine-tuning pipeline in greater detail.

#### 3.3.1 Synthetic Sequence Generation

At each fine-tuning step, a peptide sequence of 4–52 residues is sampled uniformly from the PDB training set used in Phase 1. Each position is then independently mutated to an ncAA with probability 0.2, drawn uniformly from the 189-ncAA support set of the differentiable hydrogen placement module (Section C.4). A cyclic or linear topology flag is assigned with probability 0.4 and 0.6, respectively. Head-to-tail peptide cyclicisation is an important property for peptide therapeutics, and AF3 natively accepts a cyclicity specification, in the form of a positional encoding, which embeds a head-to-tail proximity constraint, such that the first and last residues should be spatially close in the predicted structure. No MSA, templates, or inference-time steering are used; the model receives only the sequence with ncAAs and cyclicity specifications. Additional details are given in Appendix C.1.

#### 3.3.2 Fine-tuning With a Differentiable Reward Specifics

Each sampled sequence is passed to the pretrained model, which generates *M* = 4 conformations 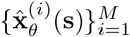 via *T* = 200 steps of diffusion sampling. These conformations are scored by SO3LR, their energies averaged, and the resulting signal used to update the model weights via gradient descent to minimize Eq. equation 2. To propagate this signal back through the model, we use the DRaFT (Directly Fine-Tuning Diffusion Models on Differentiable Rewards) framework (Clark et al., 2023), illustrated in Figure 1b and summarised in Algorithm 1.

As shown in Figure 1b and Algorithm 1, we use DRaFT-*K*, retaining the gradient graph through only the last *K* = 10 denoising steps to improve memory efficiency and reduce the risk of exploding gradients. Since SO3LR is a force field, it returns atomic forces 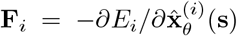 — the analytical negative gradient of the energy with respect to the final denoised coordinates — which are injected directly into the autograd graph in place of backpropagating through SO3LR itself. The full parameter gradient used in the gradient descent is then:

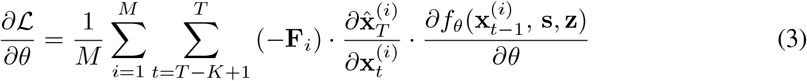

where *ℒ* is the final SO3LR energy value averaged over the *M* conformations, 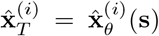, and 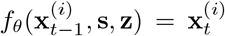, where *f*_*θ*_ denotes the diffusion model, which takes as input the atom coordinates at timestep *t −* 1 and the single **s** and pair **z** representations from the trunk, and outputs the next denoised coordinates at timestep *t*. Note that we consider the transition from 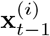 to 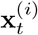 as the denoising direction.

We can decompose *∂ℒ/∂θ* into two parts: The gradient w.r.t the diffusion module weights *θ*_diff_, and the gradient w.r.t the trunk weights *θ*_trunk_. The gradient w.r.t. the diffusion weights *θ*_diff_ follows directly from Eq. 3 with *∂f*_*θ*_*/∂θ*_diff_ in place of *∂f*_*θ*_*/∂θ*. For the gradient w.r.t. the trunk weights *θ*_trunk_: the trunk runs once per training step to produce (**s, z**), which are shared across all *K* retained denoising steps and all *M* conformations. The gradient w.r.t. (**s, z**) therefore accumulates from all *K × M* calls:

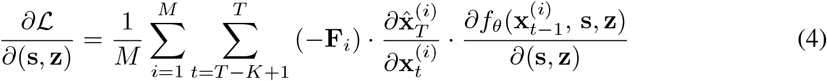

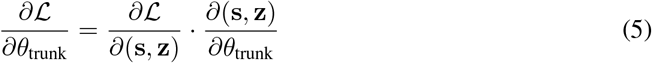

#### 3.3.3 Differentiable Hydrogen Placement

The gradient chain in Eq. 3 terminates at the heavy-atom coordinates 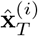 predicted by the diffusion model. However, SO3LR is an all-atom force field - it requires hydrogen atom positions, which the diffusion model does not predict. Standard structure protonation tools (e.g., RDKit, OpenBabel, OpenMM) are all either non-differentiable, and/or are parametrized for only the 20 canonical amino acids. We therefore implement a differentiable hydrogen placement module *h* in PyTorch, inspired by the approach of differentiable hydrogen placement for canonical amino acids in SideChainNet King & Koes (2021), and extend it to a set of 189 ncAAs such that 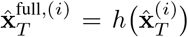 is a differentiable function of the heavy-atom coordinates. The 189 supported residues were selected by first identifying the most frequently occurring non-canonical residues across protein chains in the PDB, then incrementally expanding the set as new ncAA types were encountered during subsequent test set curation.

The placement is split into a one-time preprocessing step and a differentiable runtime step. In preprocessing, the hydrogens that need to be placed for a given residue are given by the CCD Westbrook et al. (2014). Each hydrogen to be placed is assigned a *placement rule*: a set of reference heavy atoms and a closed-form geometric construction chosen according to the local bonding environment of its parent atom (e.g., hydrogens can assume any of the geometries which constrains their placement: methyl, methylene, tetrahedral, trigonal, etc). We define placement rules for each of the 20 canonical and 189 non-canonical residues, hard-coding how each hydrogen in that residue must be positioned via differentiable operations. At runtime, hydrogens are placed according to the rules specified for their residue. We note that this hard-coding inherently limits the number of ncAAs that the model can train on, so we aim to focus efforts on making the hydrogen placement generalizable in the future. The full construction rules for each of the hydrogen placement geometries are given in Appendix C.4.

This extends the full gradient chain from Eq. 3 to:

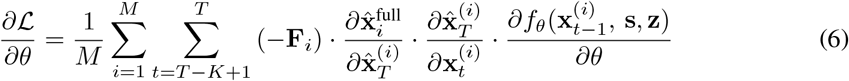

where 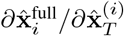 is the Jacobian of the hydrogen placement module-connecting the all-atom energy gradient (*−***F**_*i*_) to the heavy-atom output of the diffusion model.

## 4 Experiments

### 4.1 Test Set Curation

There is currently no established benchmark for ncAA peptide structure prediction. We therefore curate our own evaluation set from structures deposited in the PDB with release dates after January 1, 2024, ensuring temporal separation from our training cutoff of June 1, 2023 and our validation window (June 1, 2023–January 1, 2024).

#### Non-canonical peptide set (34 peptides)

We downloaded 45,579 structures from the RCSB and extracted all protein chains of length 4–52 residues. Sequences are then subsequently filtered via the following pipeline: (*i*) removal of sequences that are similar to the training set via MMseqs2 (Steinegger & Söding, 2017)-any chain sharing *≥* 40% sequence identity over *≥* 80% query coverage is excluded; (*ii*) removal of chains whose only non-standard residues are trivial terminal caps (ACE, NH2), unresolved (UNK) positions, or positions whose CCD codes are classified as non-polymer ligands. From the remaining polymer-only ncAA pool we retain 34 chains for which every non-canonical residue is supported by our differentiable H-placement module (Section C.4).

#### Canonical peptide set (24 peptides)

We additionally curate a canonical-only set which contain chains from the same post-2024 release window that contain no non-canonical residues, in order to monitor potential regression on standard amino acids after fine-tuning. The same MMseqs2 filter is applied.

### 4.2 Baselines

We compare against three baselines. **Boltz-2** (Passaro et al., 2025) is the publicly released check-point of the model our pretraining stage is based on; it serves as a measure of how well a general-purpose structure predictor handles ncAA chemistry without any ncAA-specific adaptation. **Protenix-v1** (Zhang et al., 2026b), **(Team et al**., **2025), Zhang et al. (2026a) is a state-of-the-art open-source AlphaFold-3-style predictor; 5 of the 34 ncAA test peptides contain residues not in Protenix’s CCD vocabulary and are excluded from its evaluation (*n*=29). Pretrained (Neopep Phase 1)** is our own model after Phase 1 pretraining only, before any RL fine-tuning.

All models run in single-sequence mode (no MSA). We generate 25 structure predictions per test peptide for all models.

### 4.3 Evaluation Metrics

We report the C*α* RMSD (**CA-RMSD**) after superposition of the predicted chain onto the ground-truth. Because our fine-tuning objective minimises a force-field energy rather than a direct structural loss, the model can produce a diverse ensemble of low-energy conformers that may not cluster tightly around the crystal structure. We therefore report the **minimum CA-RMSD** over 25 predictions (the closest the model can get to the ground truth) and additionally the **average CA-RMSD**. We also report the minimum heavy-atom RMSD (**FA-RMSD**) after superposition.

### 4.4 Results

Table 1 summarises the overall CA-RMSD averaged across each test subset. Per-peptide break-downs are provided in Appendix D.1.

**Table 1:** CA-RMSD (Å ) averaged over all test peptides (mean *±* std across peptides, min CA-RMSD is computed over the average of all the min CA-RMSDs across the test peptides). Bold indicates the best value in each column. Protenix evaluated on *n*=29 non-canonical peptides (5 excluded due to missing CCD support); all other models evaluated on *n*=34.

| Model | Non-canonical ($n=34$) | | Canonical ($n=24$) | |
| --- | --- | --- | --- | --- |
|  | Min CA-RMSD | Avg CA-RMSD | Min CA-RMSD | Avg CA-RMSD |
| Boltz-2 | 4.448 ± 1.595 | 9.047 ± 5.522 | 2.744 ± 2.605 | 3.438 ± 2.955 |
| Protenix-v1<sup>†</sup> | 2.258 ± 1.320 | 2.845 ± 1.437 | 3.005 ± 3.022 | 3.738 ± 3.039 |
| Neopep Pretrained | 2.271 ± 0.837 | 3.236 ± 0.958 | 3.046 ± 2.765 | 3.847 ± 3.105 |
| Neopep Finetuned | **1.565** ± 0.957 | **2.546** ± 1.160 | **2.211** ± 1.927 | 3.768 ± 3.078 |

<sup>†</sup>Evaluated on $n=29$ non-canonical peptides.

#### Non-canonical peptides

Neopep fine-tuned achieves the lowest average Min CA-RMSD (1.565 Å ) among all evaluated models. The large gap between Boltz-2 (4.448 Å ) and the pre-trained Neopep model (2.271 Å ) suggests that the pretraining strategy of training on PDB fragments is beneficial for establising a good structural prior for peptides. The further improvement from the energy fine-tuned model confirms that the SO3LR energy reward steers the diffusion model towards physically plausible conformations of ncAA-containing peptides. The same ordering holds for heavy-atom FA-RMSD (Table 2): Neopep fine-tuned achieves a Min FA-RMSD of 3.194 Å, compared to 3.735 Å for the pretrained model, 3.563 Å for Protenix-v1, and 5.818 Å for Boltz-2 on the non-canonical test set. Additionally, figure 2 shows the SO3LR computed average energy per atom over 25 predictions per peptide, for the non-canoncial and canoncial test sets, across the same models. We highlight the performance of our finetuned model in that it is able to predict more lower energy conformers, while also matching the eV/atom of the ground truth structrures. This is essentially a sanity check, to ensure that our fine-tuning objective worked as intended. We report statistical tests in Appendix E

**Table 2:** Minimum heavy-atom RMSD (FA-RMSD, Å ) averaged over all test peptides (mean *±* std across peptides). Bold indicates the best value in each column. Protenix evaluated on *n*=29 non-canonical peptides; all other models on *n*=34.

| Model | Min FA-RMSD, Non-canonical ($n=34$) | Min FA-RMSD, Canonical ($n=24$) |
| --- | --- | --- |
| Boltz-2 | $5.818 \pm 1.458$ | $4.035 \pm 2.550$ |
| Protenix-v1<sup>†</sup> | $3.563 \pm 1.553$ | $4.331 \pm 2.922$ |
| Neopep Pretrained | $3.735 \pm 0.804$ | $4.037 \pm 2.800$ |
| Neopep Finetuned | **$3.194 \pm 0.944$** | **$3.456 \pm 1.917$** |

<sup>†</sup>Evaluated on $n=29$ non-canonical peptides.

**Figure 2:**
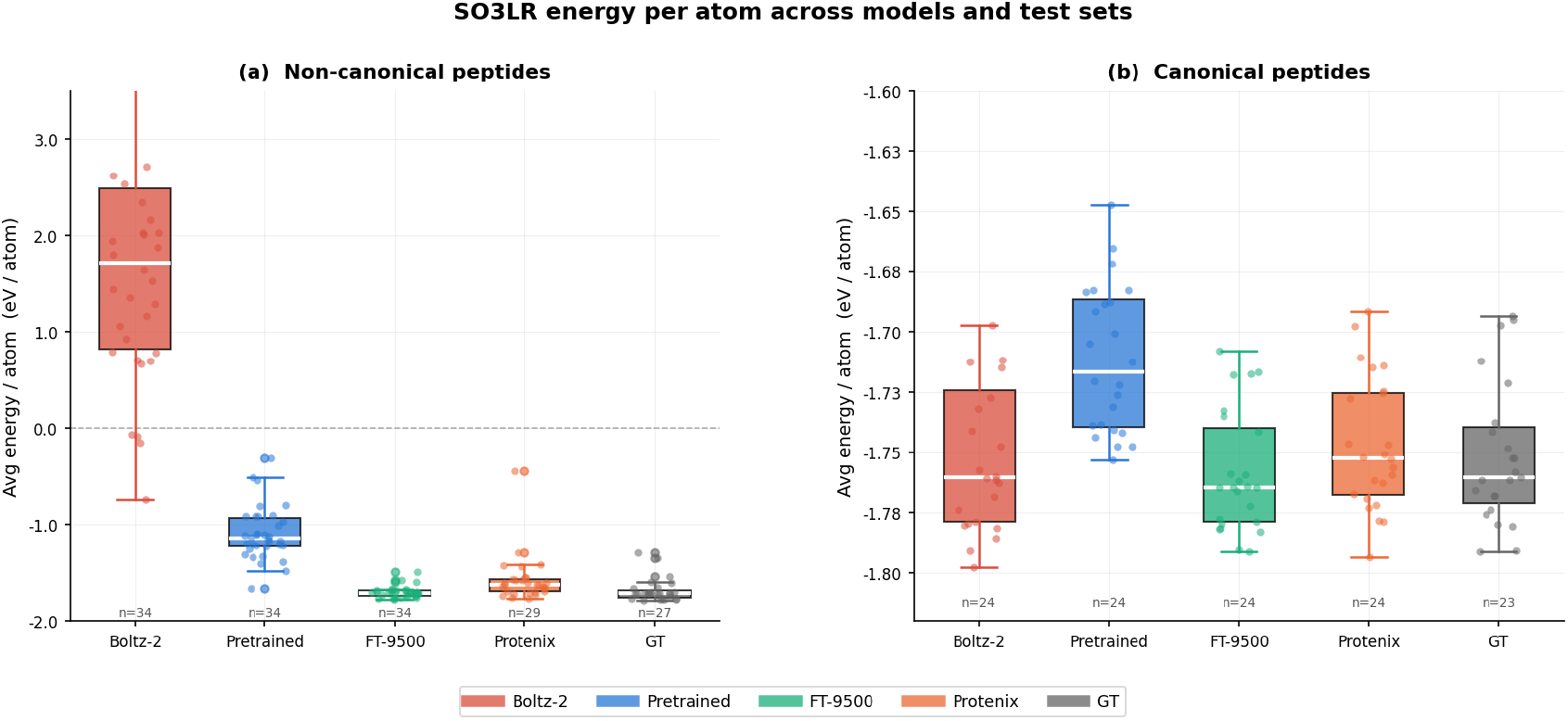
Average SO3LR energy per atom (eV/atom) over 25 predictions per peptide, across all peptides for the non-canonical (left) and canonical (right) test sets. GT = crystal structure energy. Some atoms in the ground truth for some of the test peptides were missing, so for a fair comparison of the energies, we only include ground truth peptides that have the same number of atoms as the predictions (i.e., everything is fully resolved).

#### Canonical peptides

Fine-tuning does not degrade performance on canonical sequences: Neopep fine-tuned achieves the best average Min CA-RMSD on the canonical set (2.211 Å ), outperforming both Boltz-2 (2.744 Å ) and Protenix-v1 (3.005 Å ). Neopep fine-tuned achieves a Min FA-RMSD of 3.456 Å, compared to 4.035 Å for Boltz-2, 4.037 Å for the pretrained model, and 4.331 Å for Protenix-v1 (Table 2). Additionally, we look at the SO3LR computed average energy per atom over 25 predictions per peptide in figure 2. Similar to the results for the non-canonical test set, our finetuned model outputs structures that are either matching in energy to the respective ground truth, or are slightly better. Again, we note that this is essentially a sanity check to ensure that our training objective works as intended. We report statistical tests in Appendix E

### 4.5 Case Studies

To qualitatively illustrate the effect of MLFF energy-guided fine-tuning, we examine two non-canonical peptides from the test set where the pretrained and fine-tuned models produce noticeably different structures. For each peptide we show the minimum-RMSD prediction from each model overlaid on the crystal structure, rendered with PyMOL (Schrödinger, LLC).

As shown in Figures 3, and 4 the pretrained-only model produces a conformation that appears to not be aligned well to the ground truth, and pymol was unable to infer some bonds, which is why the structure looks like it has “breaks” in it. We then examine a structure after energy tuning, and observe that it appears to be fully connected, which means that pymol correctly inferred bonds from atomic positions, which is reflected in the very low CA-RMSD.

**Figure 3:**
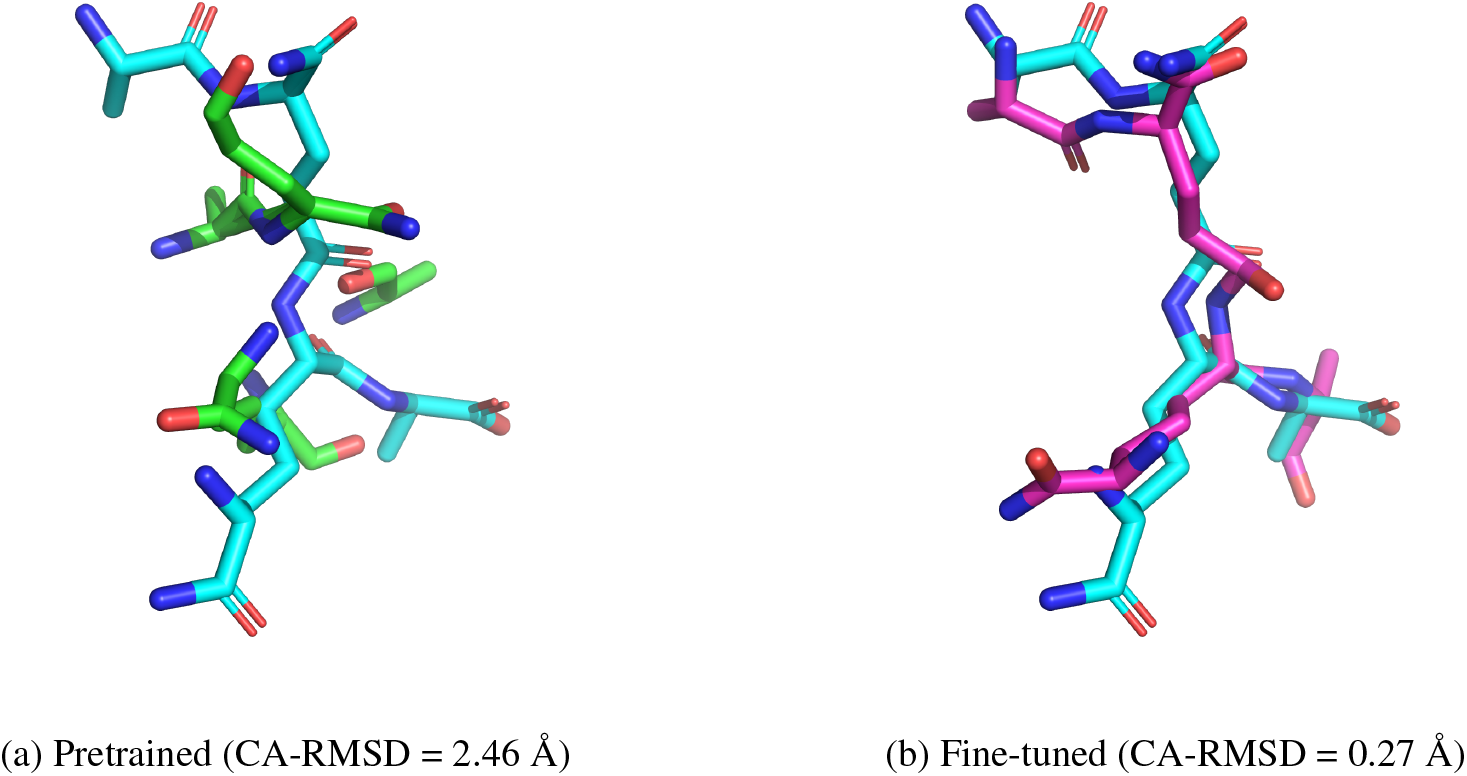
**8pxy B** - predicted structure (green, pink) overlaid on the ground truth structure (blue). Left: Neopep pretrained model. Right: Neopep fine-tuned (FT-9500). Fine-tuning corrects ‘breaks’ in structures, allowing pymol to better infer bonds from better placed atoms.

**Figure 4:**
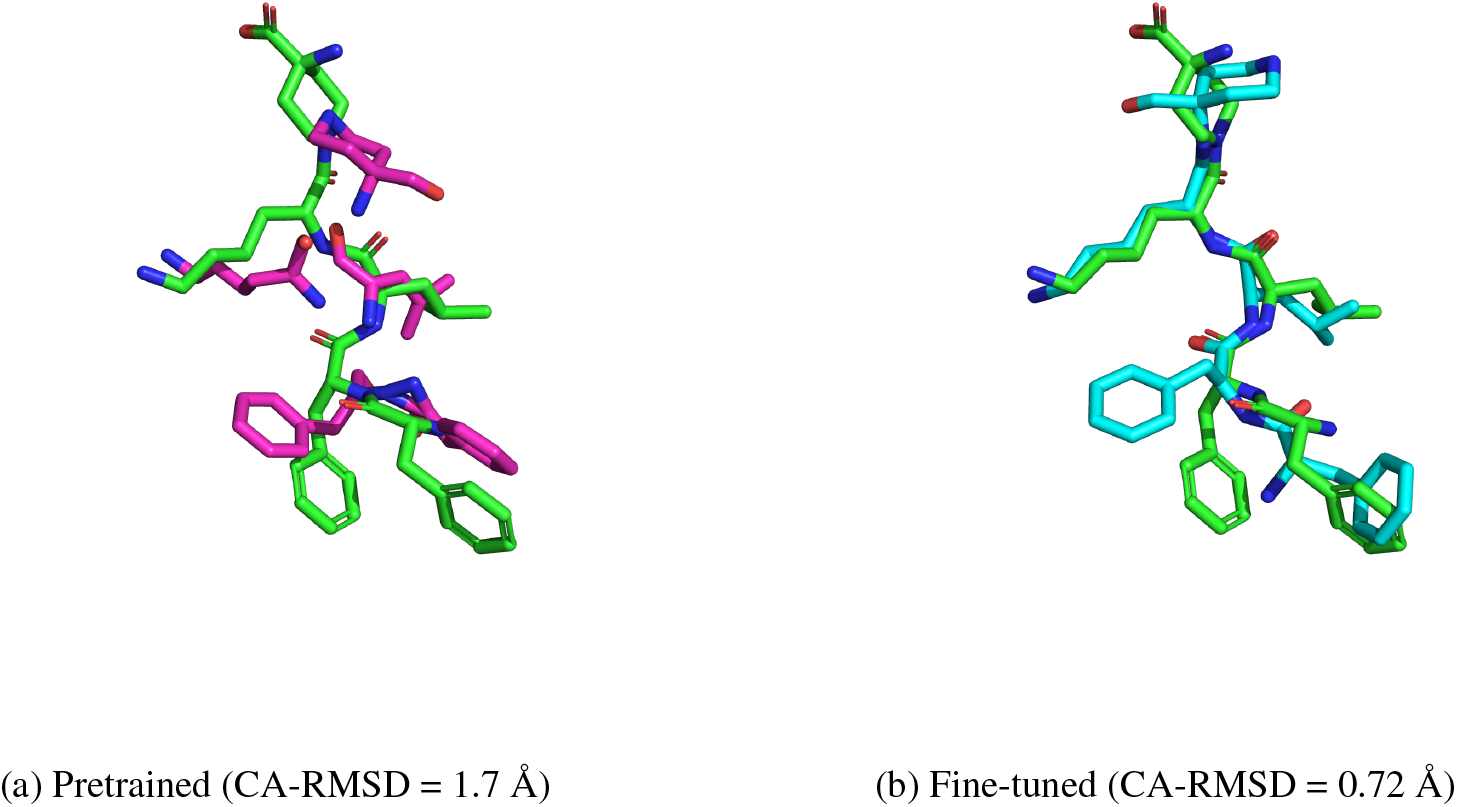
**22es F** - predicted structure (pink, blue) overlaid on the ground truth structure. Left: Neopep pretrained model. Right: Neopep fine-tuned (FT-9500). Fine-tuning corrects ‘breaks’ in structure, allowing pymol to better infer bonds from better placed atoms.

## AI use statement

In this work, we used generative AI tools to aid in the writing and planning of the paper contents, especially getting lateX-style equations. AI was also use to write code scripts to automate processes. For instance, a script to search through a database, or a script to write hydrogen placement functions for each of our parametrized 189 ncAAs given a list of rules to follow. We do not however, use AI to prove any sort of mathematical claims or for the research ideation process. Wherever we did use LLM assistance (when writing the paper, for isntance) we made sure to thoroughly check its outputs and understand everything before deciding to include it. We take responsibility for the final content of this work, including text, claims or artifacts produced with the aid of generative AI.

## Ethics statement

Our models leverage already-public data for training, such as the Protein Databank. As such, we feel that we do not have any instances where the paper submission raises questions in the code of Ethics.

## Reproducibility statement

We provide experiments in section 4, detailing how we curated our datasets and which baselines we used. The appendix has additional details on our augmentations to the Boltz2/AF3 architecture, pretraining, and finetuning, as well as ablations.

## Acknowledgments

We thank the authors of SO3LR: Professor Alexandre Tkatchenko and Adil Kabylda (University of Luxembourg), for their generous insights and advice on the direction of this project.

## A Appendix

## A Architecture Details

### A.1 Model Configuration

Table 3 summarizes the key hyperparameters of the Neopep model. We follow the AF3/Boltz architecture but reduce the model size relative to the full Boltz-2 configuration to allow more efficient training.

**Table 3:**
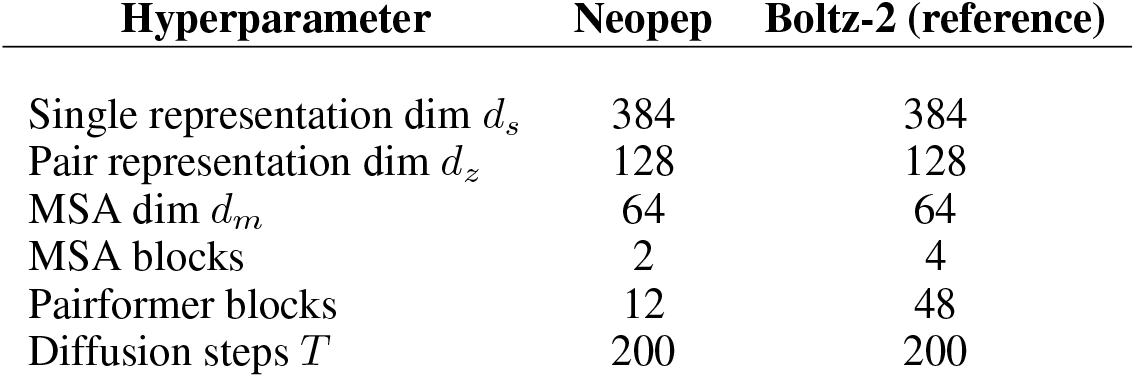
Neopep model hyperparameters.

| Hyperparameter | Neopep | Boltz-2 (reference) |
| --- | --- | --- |
| Single representation dim $d_s$ | 384 | 384 |
| Pair representation dim $d_z$ | 128 | 128 |
| MSA dim $d_m$ | 64 | 64 |
| MSA blocks | 2 | 4 |
| Pairformer blocks | 12 | 48 |
| Diffusion steps $T$ | 200 | 200 |

**Table 4:** Pretraining hyperparameters.

| Hyperparameter | Value |
| --- | --- |
| Effective batch size | 128 |
| Peak learning rate | $1.8 \times 10^{-3}$ |
| LR schedule | AF3: linear warmup 1,000 steps,<br>decay $\times 0.95$ every 50,000 steps<br>after step 50,000 |
| Training steps | $\sim 8,400$ optimizer steps |
| GPUs | 4 |
| GPU type | H100 |

**Table 5:** Fine-tuning hyperparameters.

| Hyperparameter | Value |
| --- | --- |
| Optimizer | AdamW |
| Learning rate | $10^{-5}$ |
| Weight decay | 0.1 |
| $(\beta_1, \beta_2)$ | (0.9, 0.999) |
| Gradient clip (norm) | 1.0 |
| DRaFT-$K$ | 10 |
| Diffusion steps $T$ | 200 |
| Conformations $M$ | 4 |
| Trunk recycles $N_r$ | 3 |
| Random seed | 42 |
| GPU | H100 (1×) |
| Training Steps | 9500 steps |

#### MSA module

We include a lightweight MSA module with *d*_*m*_=64 and 2 MSA blocks (vs. 4 in Boltz-2). During pretraining, the MSA module processes up to `max_seqs`= 512 sequences. During fine-tuning, the MSA module is disabled (`max_seqs`= 0), and the model receives only the single input sequence with no alignment.

#### Template module

We disable the template module entirely (`use_templates=False`). No template embedder is instantiated and no template features are computed at any stage of training or inference.

#### Confidence and affinity modules

The confidence and affinity head modules present in Boltz-2 are not used. These heads were removed to reduce memory footprint and because our training objective does not require predicted confidence scores or affinity.

## B Pretraining Details

### B.1 Dataset

Our pretraining dataset is drawn from the Protein Data Bank (PDB), following the same structure processing and filtering methods outlined in the Boltz source code and paper Passaro et al. (2025). Specifically, we use a training cutoff date of June 1, 2023, leaving structures deposited after this date for our validation and test sets.

Because our target application is peptides (3–50 residues), we train on *protein fragments* of length 3–50 residues sampled from longer PDB chains. Each fragment is sampled by choosing a contiguous window of the desired length uniformly at random from a sampled protein chain, ensuring the window does not cross chain breaks. This allows us to make use of all proteins in the PDB. MSA coverage is reduced relative to full proteins; we therefore cap the number of MSA sequences at 512 and disable template search entirely. All other data processing, clustering, and sampling follow Passaro et al. (2025). Figure 5 shows the diffusion loss curves during pre-training.

**Figure 5:**
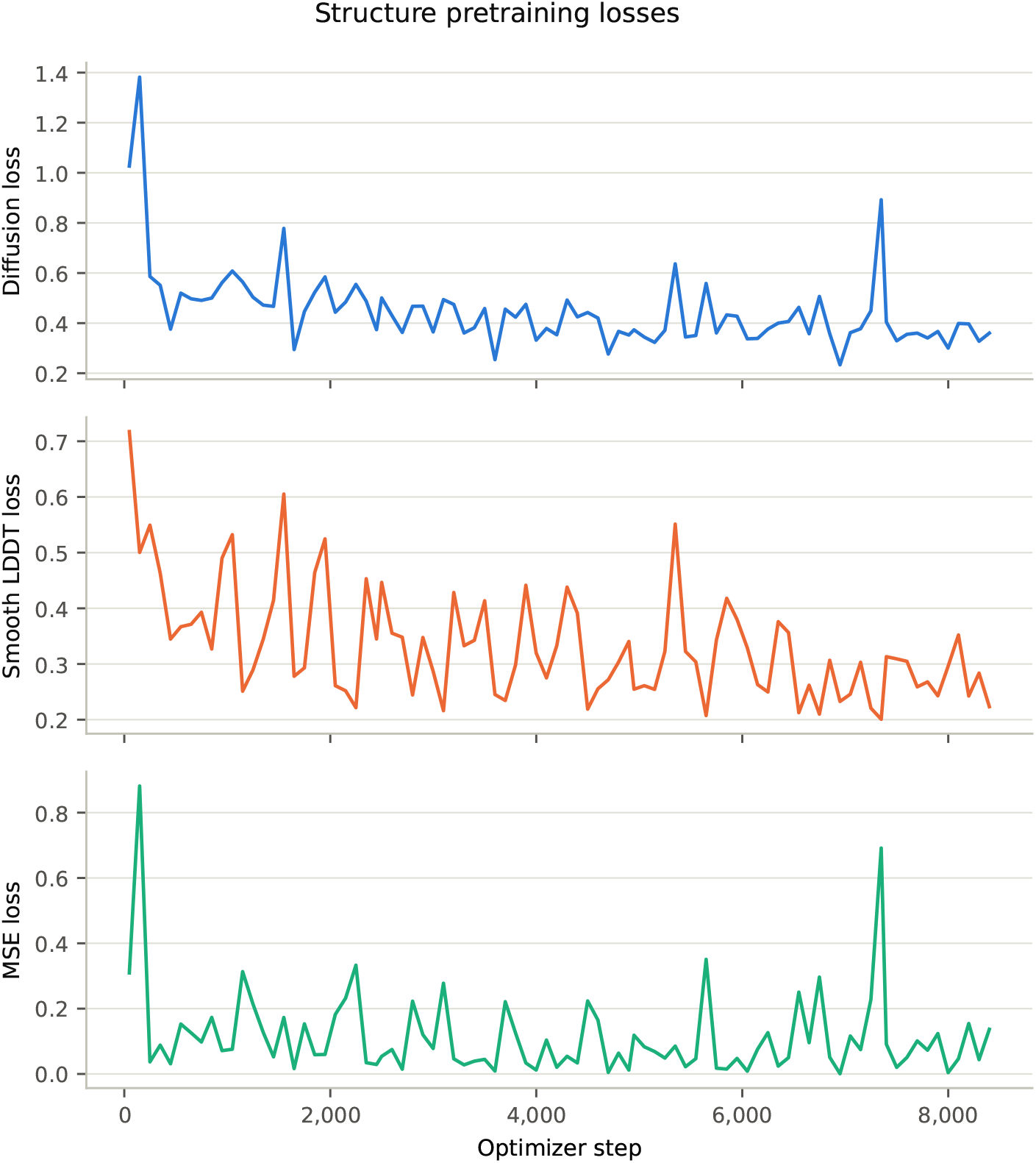
Training losses over ∼8,400 optimizer steps of structure pretraining. **Top:** diffusion loss. **Middle:** smooth LDDT loss. **Bottom:** coordinate MSE loss.

### B.2 Training Configuration

Training follows the AF3 scheme with no concept of epochs; the training distribution is sampled continuously.

## C Fine-tuning Details

### C.1 Synthetic Sequence Generation

At each fine-tuning step, a peptide sequence is drawn from the PDB training set used in Phase 1 via the following procedure:

1. **Chain selection**. A single protein chain of length 4–52 residues is sampled uniformly from a pre-filtered chain index built from the training manifest. Each entry in the index is a (PDB id, chain idx) pair that has already passed the length filter, so every draw is guaranteed valid without rejection sampling.
2. **Cap stripping**. Terminal capping residues (N-terminal ACE, C-terminal NH2) are removed before any further processing.
3. **ncAA substitution**. Non-standard residues already present in the sampled chain that are not in the 189-ncAA support set are replaced with a residue drawn uniformly from the support set. Each remaining position (whether a canonical or already-supported ncAA) is then independently mutated to a uniformly drawn ncAA with probability *p*_ncAA_ = 0.2.
4. **Cyclicity assignment**. The peptide is designated cyclic with probability *p*_cyc_ = 0.4, and linear with probability 0.6.
5. **YAML encoding**. The resulting residue list and cyclicity flag are encoded into a Boltz-compatible YAML input. Canonical residues are encoded by their one-letter code; ncAAs are encoded as CCD three-letter codes in the `modifications` field. The MSA field is set to `empty`, so no alignment is used.

The 189 supported ncAAs are defined by the set `SUPPORTED_NCAAS` in the differentiable hydrogen placement module (Section C.4).

### C.2 Optimization Hyperparameters

The pretrained checkpoint is loaded and the model is fine-tuned end-to-end on both the diffusion and trunk weights, with SO3LR frozen throughout; each step processes one sequence with *M* =4 conformations.

#### Energy pooling

We use average pooling over the *M* conformations as the default. We also evaluate a max-pooling variant, 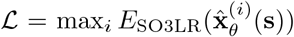, which focuses the gradient on the worst-energy conformation; results are reported in Section D.

#### SO3LR inputs and charge computation

SO3LR (Kabylda et al., 2025) is an equivariant machine-learned force field that takes as input an *ASE* `Atoms` object specifying atomic positions, element symbols, and a scalar net charge, and returns the potential energy *E* (in eV) together with atomic forces **F** = *− ∂E/∂***x** (in eV/Å ). Because SO3LR is charge-aware, we compute an approximate net integer charge for each peptide at pH 7.4 using a lookup table of side-chain formal charges ( `_RESIDUE_CHARGES` in the differentiable H-placement module). Only charged side-chain groups are accounted for. The resulting integer charge is passed to SO3LR via `atoms.info[“charge”]` at each forward call. SO3LR is kept frozen throughout all fine-tuning; no parameters of the force field are updated.

### C.3 Supported Non-Canonical Amino Acids

The differentiable hydrogen placement module is parametrised for the following 189 ncAAs (CCD three-letter codes). Only residues in this set may appear in fine-tuning sequences; canonical amino acids are always supported in addition. The 189 supported ncAA types were assembled in two stages. We first identified the most frequently occurring non-canonical residues across protein chains in the PDB and hand-specified differentiable H-placement rules for those residues. The set was then expanded incrementally as new ncAA types were encountered during curation of our test set.

A practical implication is that extending Neopep to new chemistries is straightforward: adding a new ncAA requires only writing one additional placement rule (specifying the hydrogen geometry for that residue type), and running further RL fine-tuning steps with the expanded residue pool. The 189-residue set listed below should therefore be seen as a starting point rather than a limit of the approach.

~~~
 004, 00E, 0A1, 0W6, 28J, 2TL, 33X, 3EG, 3FG, 4HH, 4J2, 4WQ, 5CW, 66E,
6ZS, 7C9, 7T2, A1A5S, A1A5T, A1A8T, A1AL8, A1AMB, A1AMC, A1AT6, A1AT7,
 A1B6Z, A1BC0, A1CCQ, A1CCR, A1CCS, A1CHA, A1CLH, A1CZ7, A1D5E, A1D5P,
 A1EAK, A1ECM, A1EQM, A1EUR, A1I4F, A1I67, A1I8Q, A1I8R, A1ICB, A1ICC,
 A1IFI, A1IJP, A1IKE, A1IUO, A1J0G, A1J16, A1JND, A1L3I, A1L4D, A1L7K,
A1L9X, A1MFG, ABA, AC5, ACA, AIB, ALO, ALY, B3S, BAL, BB8, BMT, CGU, CHG,
 CSD, CSO, CSS, CSX, D4P, DAB, DAL, DAM, DAR, DAS, DGL, DGN, DI8, DIL,
 DLE, DLY, DPN, DPP, DPR, DSG, DSN, DTH, DTR, DTY, DVA, EEP, FME, G5G,
 GHP, HIC, HJV, HPE, HRG, HY3, HYP, IAE, IAS, IC0, IIL, IML, JGO, JXR,
 JYC, KBE, KCR, KJW, LE1, LTU, M70, MAA, MEA, MHS, MLE, MLU, MLY, MLZ,
 MVA, MYN, N2C, N9P, NAL, NCY, NH2, NIY, NLE, NVA, NZC, O7G, OCS, OIC,
 OMY, OMZ, ORD, ORN, OUK, OUR, PCA, PHA, PIV, PPN, PQ4, PTR, QDD, QOL,
 QQB, QUJ, QUK, QVE, QVS, SAC, SAR, SEP, SFE, SLL, SNC, SOQ, TBG, TIH,
 TPO, TY5, TYS, UAL, UKD, UN1, URL, UX8, WEH, WFP, WLU, WPA, WVL, WWB,
 X60, XRW, YC3, YNM, YYA, ZAE, ZGL, ZY9
~~~

### C.4 Hydrogen Placement Geometry Rules

The differentiable hydrogen placement module, takes inspiration from the differentiable hydrogen adder in King & Koes (2021), and assigns each hydrogen atom a *placement rule* determined by its local bonding environment. Six rules are defined; all operations are implemented in PyTorch and are fully differentiable with respect to the heavy-atom coordinates. In each rule, *C* denotes the heavy parent atom, *R*_1_, *R*_2_, … denote its bonded reference heavy atoms, *d* is the target C–H bond length, 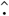 denotes *l*_2_ normalisation, and *×* denotes the cross product.

#### Rule 1: Bisector (sp^2^, 2 reference atoms)

Used for aromatic, carbonyl, and planar amine hydrogens. The hydrogen lies in the plane spanned by *R*_1_–*C*–*R*_2_, along the bisector of the supplementary angle:

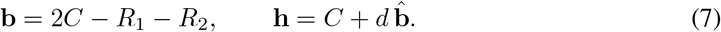

#### Rule 2: Methine (sp^3^, 2 reference atoms)

Used for CH groups where the hydrogen lies opposite the bisector of the *R*_1_–*C*–*R*_2_ angle, placing it in a roughly tetrahedral geometry. The construction is identical to Rule 1:

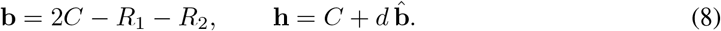

The distinction from the bisector rule is purely in which bonding environment triggers it (sp^3^ single-bond context vs. sp^2^ double-bond context).

#### Rule 3: Tetrahedral fourth (sp^3^ 4th, 3 reference atoms)

Used for CH groups in a tetrahedral environment with 3 known substituents. The hydrogen is placed opposite the centroid of the three reference atoms:

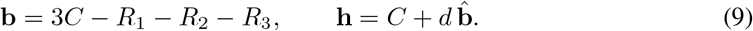

#### Rule 4: Methylene (geminal pair, 2 reference atoms)

Used for CH_2_ groups. A local orthonormal frame is constructed from the bisector direction **u** and the plane normal **v**:

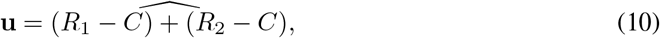

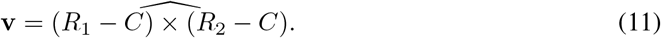

The two geminal hydrogens are placed symmetrically about the *C*–**u** axis:

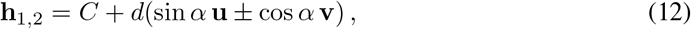

where *α* is the half-opening angle derived from the ideal H–C–H bond angle for the local geometry.

#### Rule 5: Methyl (cone, 3 hydrogens)

Used for CH_3_ groups. Three hydrogens are placed on a cone of half-angle *θ* around the *C*–*R*_1_ bond axis. An orthonormal frame (**a, m, n**) is constructed with 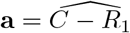 as the axis:

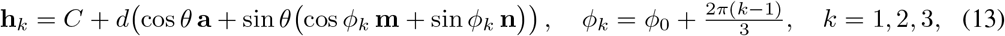

where *ϕ*_0_ is determined by the dihedral of the first hydrogen relative to a reference atom on *R*_1_, and *θ* is the ideal tetrahedral cone half-angle.

#### Rule 6: NERF

The Natural Extension Reference Frame (NERF) rule handles hydrogens whose placement requires a dihedral angle *ϕ* relative to a third reference atom *A*, in addition to the bond length *d* and bond angle *θ*. Given ordered reference atoms *A, B, C* (with the hydrogen bonded to *C*), an orthonormal local frame is constructed:

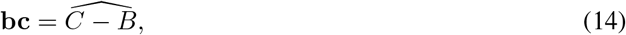

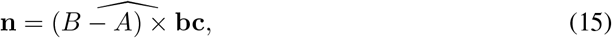

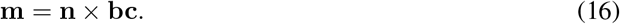

The hydrogen position is then:

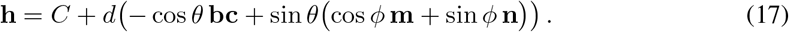

All parameters (*d, θ, ϕ*) are precomputed from the CCD geometry at preprocessing time and stored as a `HPlacementSpec` dataclass.

## D Ablations

### Number of conformations *M*

At each fine-tuning step, *M* heavy-atom conformations are sampled from the diffusion model, protonated, and scored by SO3LR; their energies are averaged to form the reward signal. Larger *M×* reduces gradient variance at the cost of *M* more MLFF evaluations per step.

Figure 6 shows energy per atom over 1,200 training steps for *M ∈ {*4, 8, 12, 16}. All four settings converge to approximately *−* 1.50 to *−* 1.54 eV/atom, with no statistically meaningful difference in final energy. The rolling means are nearly indistinguishable after step *∼* 600, indicating that the variance reduction from larger *M* does not translate into a better optimum within this training budget. We adopt *M* = 4 as the default.

**Figure 6:**
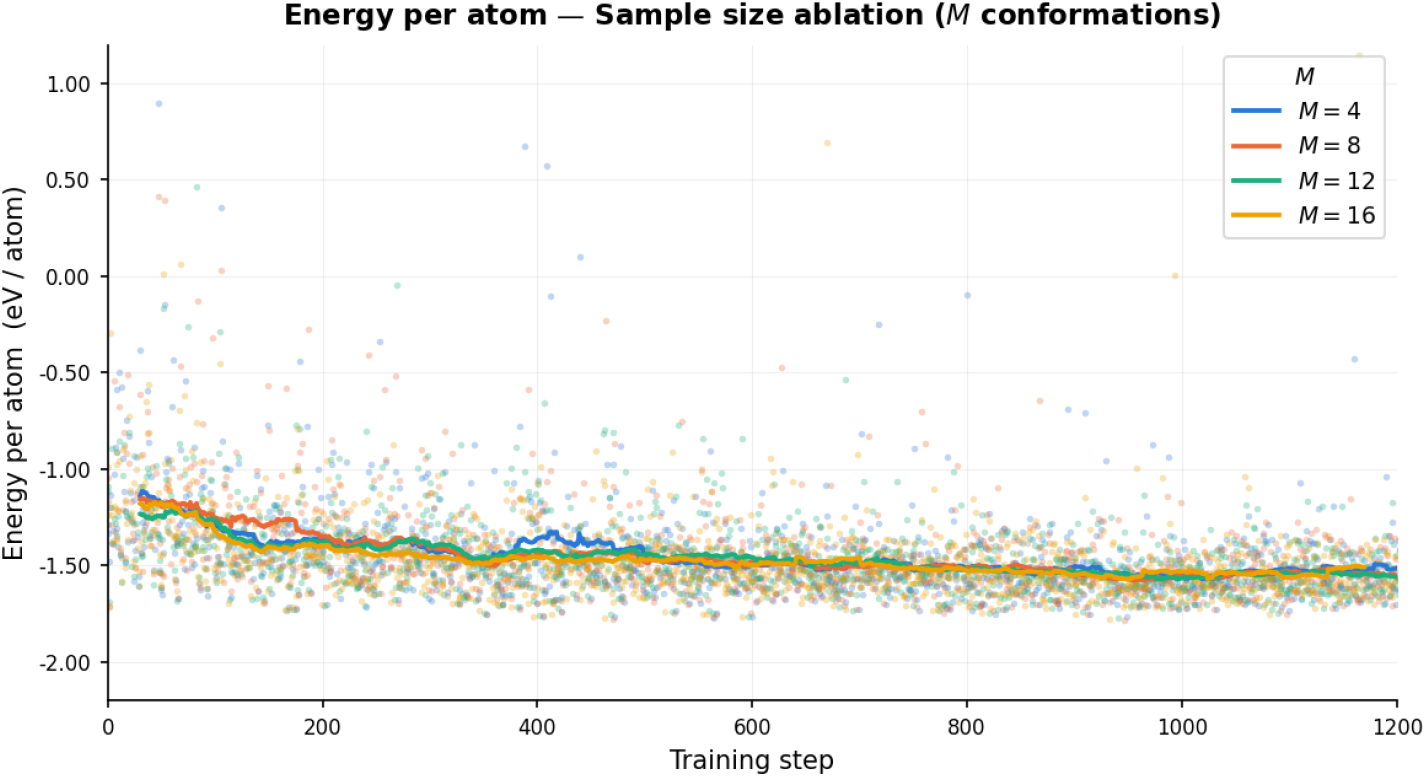
Energy per atom (eV/atom, lower is better) during RL fine-tuning for *M ∈ {*4, 8, 12, 16} sampled conformations. Dots show per-step values; solid lines show a 60-step rolling mean. All variants converge to the same final energy, motivating the choice of *M* = 4.

### Energy pooling: average vs. maximum

During RL fine-tuning, *M* = 4 conformations are sampled per peptide and scored by SO3LR; the per-conformation energies must be aggregated into a single reward signal for the gradient update. We compare two strategies: average pooling 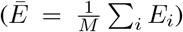 and max-pooling (*Ē* = max_*i*_ *E*_*i*_), which focuses the gradient on the highest-energy (least stable) conformation found.

Figure 7 shows the energy per atom (eV/atom) over [1,200] fine-tuning steps for both variants. Both methods drive the mean energy from approximately *−* 1.1 eV/atom to *−* 1.5 eV/atom over the course of training. The two strategies converge to similar final energies, with virtually no difference in training performance. We adopt average pooling as the default.

**Figure 7:**
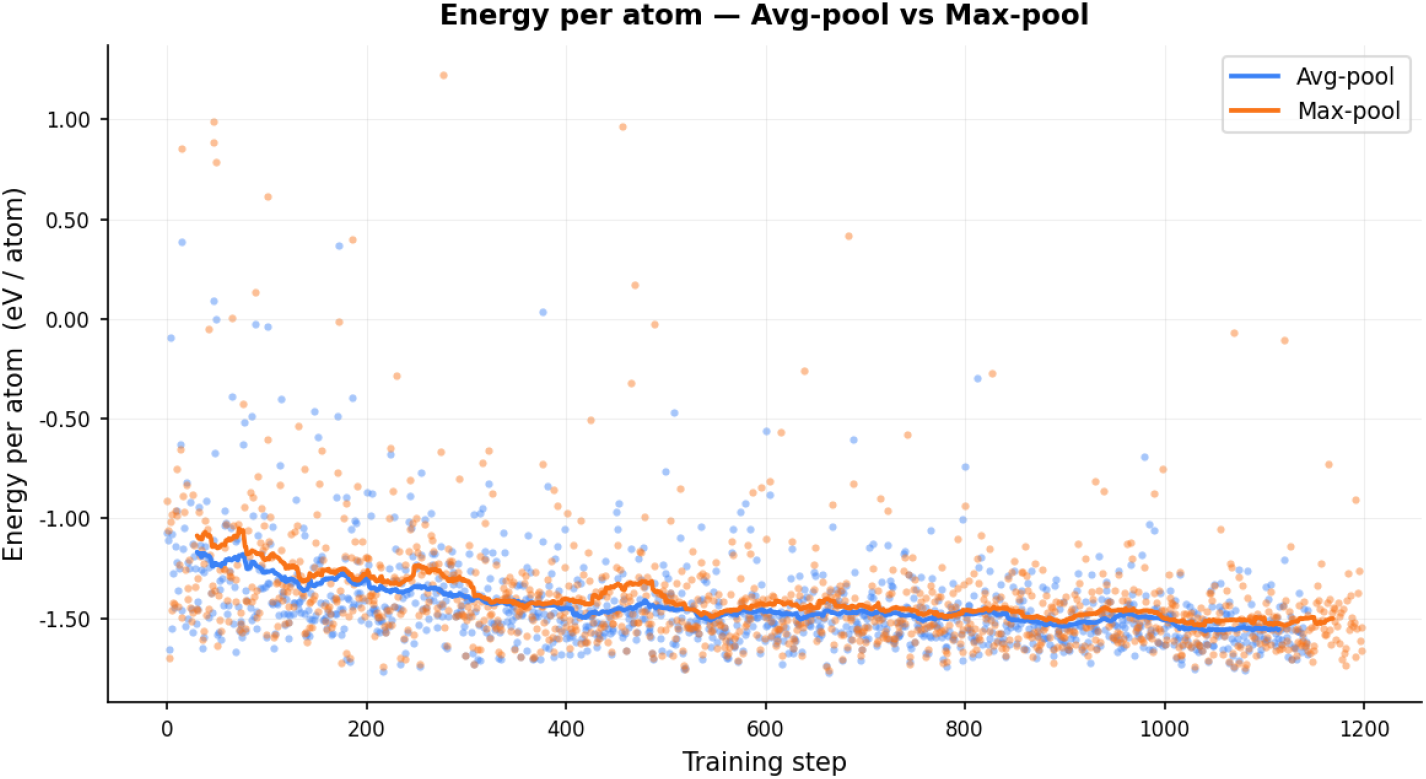
Energy per atom (eV/atom, lower is better) during RL fine-tuning for average pooling (blue) and max-pooling (orange) over *M* = 4 sampled conformations. Dots show per-step values; solid lines show a rolling mean.

### D.1 Per-Peptide Results: Non-Canonical Test Set

Table 6 reports per-peptide minimum CA-RMSD, minimum FA-RMSD, and average CA-RMSD (mean *±* std over 25 predictions) for all 34 non-canonical test peptides. Dashes indicate peptides not supported by Protenix. Bold marks the best (lowest) value in each metric row.

**Table 6:**
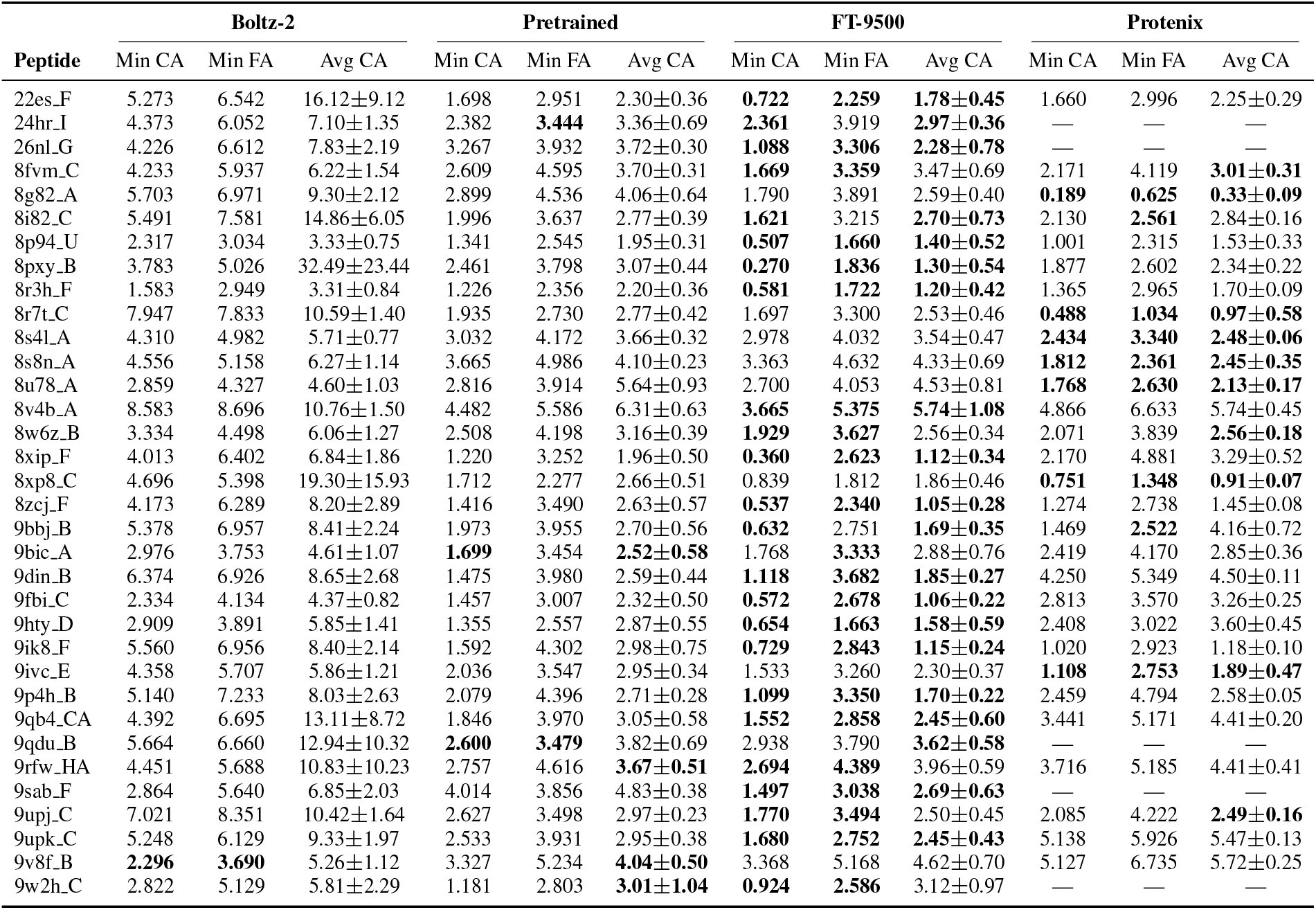
Per-peptide RMSD results (Å ) for the non-canonical test set (*n*=34). Min = best of 25 predictions; Avg = mean *±* SD over 25 predictions. Bold marks the best (lowest) value for each metric within a row. Protenix was evaluated on *n*=29 (5 peptides marked — not supported).

## E Statistical Tests

### E.1 SO3LR Energy Comparisons

Table 7 reports paired Wilcoxon signed-rank tests on mean SO3LR energy per atom. Predictions from FT-9500 (Neopep finetuned) attains lower energy than all three baselines on both test sets (canonical and non-canonical).

**Table 7:** Paired Wilcoxon signed-rank tests on mean SO3LR energy per atom (for each test peptide, we take the mean over the 25 energy per atom values), testing *E*(FT-9500) *< E*(baseline). One-sided, Holm-corrected across the three baselines within each test set. ^*\**^ denotes *p <* 0.05. *r*_rb_ is the matched-pairs rank-biserial correlation, where *−* 1 indicates FT-9500 attains lower energy on every peptide. Protenix does not support 5 non-canonical peptides, so each comparison uses pairwise complete cases; restricting all comparisons to the 29 peptides covered by every model gives identical conclusions (*p* = 5.6 *×* 10^*−*9^ against Boltz-2 and the pretrained model, *p* = 4.7 *×* 10^*−*8^ against Protenix).

| Test set | Baseline | $n$ | $p$ | $r_{\text{rb}}$ |
| --- | --- | --- | --- | --- |
| Non-canonical | Boltz-2 | 34 | $< 10^{-7*}$ | $-1.00$ |
| | Pretrained | 34 | $< 10^{-7*}$ | $-1.00$ |
| | Protenix | 29 | $< 10^{-7*}$ | $-0.96$ |
| Canonical | Boltz-2 | 24 | $0.008^*$ | $-0.55$ |
| | Pretrained | 24 | $1.8 \times 10^{-7*}$ | $-1.00$ |
| | Protenix | 24 | $6.0 \times 10^{-7*}$ | $-0.98$ |

### E.2 RMSD Comparisons

We apply the same procedure used for the energy comparisons to the structural metrics reported in Tables 1 and 2. For each metric, FT-9500 is compared against each baseline with a one-sided Wilcoxon signed-rank test paired by test peptide, Holm-corrected across the three baselines within each metric. Tables 8 and 9 report the statistical tests.

**Table 8:** Paired Wilcoxon signed-rank tests on the **non-canonical** test set, testing whether FT-9500 attains lower RMSD than each baseline. One-sided, Holm-corrected across the three baselines within each metric. ^*\**^ denotes *p <* 0.05. *r*_rb_ is the matched-pairs rank-biserial correlation, where *–* 1 ndicates FT-9500 is lower on every peptide. Protenix does not support 5 non-canonical peptides, so those comparisons use *n*=29.

| Metric | Baseline | $n$ | $p$ | $r_{\text{rb}}$ |
| --- | --- | --- | --- | --- |
| Min CA-RMSD | Boltz-2 | 34 | $< 10^{-7*}$ | $-0.99$ |
| | Pretrained | 34 | $< 10^{-7*}$ | $-0.94$ |
| | Protenix | 29 | $0.002^*$ | $-0.60$ |
| Avg CA-RMSD | Boltz-2 | 34 | $< 10^{-7*}$ | $-1.00$ |
| | Pretrained | 34 | $1.0 \times 10^{-6*}$ | $-0.87$ |
| | Protenix | 29 | $0.078$ | $-0.31$ |
| Min FA-RMSD | Boltz-2 | 34 | $< 10^{-7*}$ | $-0.97$ |
| | Pretrained | 34 | $2.8 \times 10^{-6*}$ | $-0.84$ |
| | Protenix | 29 | $0.030^*$ | $-0.40$ |

**Table 9:**
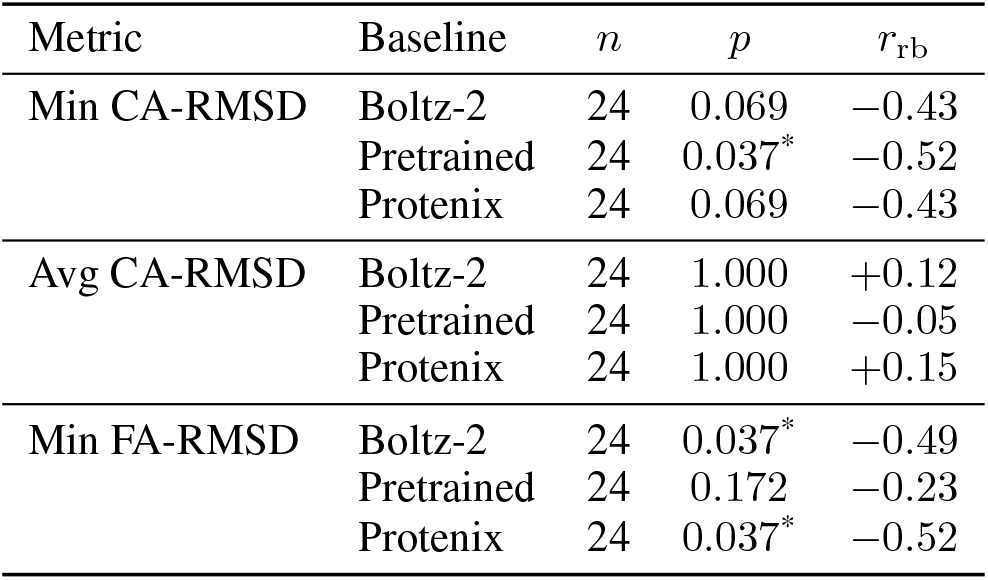
Paired Wilcoxon signed-rank tests on the canonical test set (*n*=24 for all comparisons), with the same procedure as Table 8. Positive *r*_rb_ indicates the baseline attains the lower value.

| Metric | Baseline | $n$ | $p$ | $r_{\text{rb}}$ |
| --- | --- | --- | --- | --- |
| Min CA-RMSD | Boltz-2 | 24 | $0.069$ | $-0.43$ |
| | Pretrained | 24 | $0.037^*$ | $-0.52$ |
| | Protenix | 24 | $0.069$ | $-0.43$ |
| Avg CA-RMSD | Boltz-2 | 24 | $1.000$ | $+0.12$ |
| | Pretrained | 24 | $1.000$ | $-0.05$ |
| | Protenix | 24 | $1.000$ | $+0.15$ |
| Min FA-RMSD | Boltz-2 | 24 | $0.037^*$ | $-0.49$ |
| | Pretrained | 24 | $0.172$ | $-0.23$ |
| | Protenix | 24 | $0.037^*$ | $-0.52$ |

## F Limitations

A key assumption throughout this work is the exclusion of explicit solvent. Solvation effects are well-known to influence biomolecular structure, yet incorporating them into our training scheme was infeasible for two reasons: the limited compute available to us, and the additional complexity introduced by the stochasticity of solvent configurations-a single peptide can adopt many valid solvation shells, and capturing this effect faithfully would require averaging over an ensemble of such configurations during training. We therefore restrict our structures to include hydrogen atoms and leave the incorporation of solvation effects to future work.

We also observed that the RL-based energy fine-tuning depends critically on the quality of the structural prior. When the candidate structures generated by the pretrained model are severely flawed, the RL mechanism has little to build on, and fine-tuning fails to recover a physically reasonable geometry. In other words, the energy-guided RL stage is only as strong as the prior it refines. We note that this conclusion is based on qualitative inspection of a small number of pretrained versus fine-tuned structures; we did not conduct systematic ablations or quantitative experiments to characterize this failure mode, and we leave a rigorous analysis to future work.

Lastly, since our model was trained via RL on only a fixed number of ncAAs, we cannot guarantee generalizability across the space of all possible ncAAs, though the architecture permits this in principle, as it operates on atom-level granularity. We leave future work on building a more generalizable model that can generalize to unseen ncAAs during training.

